# How Well Do Commonly Used Co-Contraction Indices Approximate Lower Limb Joint Stiffness Trends during Gait?

**DOI:** 10.1101/2020.08.05.238873

**Authors:** Geng Li, Mohammad S. Shourijeh, Di Ao, Carolynn Patten, Benjamin J. Fregly

**Author notes:** Correspondence: Benjamin J. Fregly.

## Abstract

Muscle co-contraction generates joint stiffness to improve stability and accuracy during limb movement but at the expense of higher energetic cost. The quantification of joint stiffness generated from muscle co-contraction is difficult through both experimental and computational means for its benefit and cost to be assessed. Quantification of muscle co-contraction may offer an alternative path for estimating joint stiffness. By choosing the commonly used Co-Contraction Indices (CCIs) to represent muscle co-contraction, this study investigated the feasibility of using CCI to approximate lower limb joint stiffness trends during gait. A calibrated EMG-driven musculoskeletal model of a hemiparetic individual post-stroke from a previous study was used to generate the quantities required for CCI calculation and model-based estimation of joint stiffness. A total of 14 classes of CCIs for various combinations of antagonistic muscle pairs were calculated based on two common CCI formulations, each with 7 types of quantities that included variations of electromyography (EMG) signals and joint moments from the muscles. Correlations between CCIs and model-based estimates of sagittal plane stiffness of the lower extremity joints (hip, knee, ankle) were computed. Although moderate to strong correlation was observed between some CCI formulations and the corresponding joint stiffness, these associations were highly dependent on the methodological choices made for CCI computation. The overall findings of this study were the following: (1) the formulation proposed by Rudolph et al. (2000), *CCI*_*1*_, was more correlated with joint stiffness than that of Falconer and Winter (1985); (2) Moment-based *CCI*_*1*_ from individual antagonistic muscle pairs was more correlated than EMG-based *CCI*_*1*_; (3) EMG signals with calibrated electromechanical delay and joint moment generated by individual muscle without normalization to a reference value were the most correlated for EMG-based *CCI*_*1*_ and moment-based *CCI*_*1*_, respectively. The combination of antagonistic muscle pairs for most correlated within each CCI class was also identified. By using CCI to approximate joint stiffness trends, this study may open an alternative path to studying joint stiffness.

## 1 Introduction

Muscle co-contraction refers to the simultaneous activation of muscles on opposite sides of a joint and is arguably an important mechanism used by the nervous system to regulate joint stability (Hirokawa et al., 1991; McGill et al., 2003) and provide movement accuracy (Gribble et al., 2003; Missenard et al., 2008). It is further argued that individuals who have suffered orthopedic injuries or neuromuscular disorders use elevated levels of muscle co-contraction (Lamontagne et al., 2000; Rudolph et al., 2000; Higginson et al., 2006; McGinnis et al., 2013) to generate additional joint stiffness so as to compensate for the lack of joint stability (Gollhofer et al., 1984; Kuitunen et al., 2002; Mohr et al., 2018), although evidence in support of this premise is equivocal (Banks et al., 2017). While co-contraction increases joint stiffness that may improve stability (Latash and Huang, 2015) and accuracy (Wong et al., 2009) of limb movement, it does so at the expense of increased energetic cost (Moore et al., 2014). Quantification of joint stiffness is therefore critical to the understanding of how this quantity adds both benefit and cost to dynamic joint movements such as gait. However, joint stiffness is difficult to either measure experimentally (Pfeifer et al., 2012) or calculate computationally as its determination requires appropriate interpretation of muscle recruitment strategies and muscle-tendon modeling (Sartori et al., 2015).

Quantification of muscle co-contraction may offer an alternative for estimating joint stiffness. Although muscle co-contraction and joint stiffness have been found to be related in previous studies (Kuitunen et al., 2002; McGinnis et al., 2013; Collins et al., 2014), the relationship between these two quantities remains poorly understood, as initially noted by Hortobágyi and Devita (2000). One issue is that previous studies have quantified joint stiffness primarily in the form of quasi-stiffness. Joint quasi-stiffness is described as the gradient of the torque-angle curve rather than the true characterization of joint stiffness (Rouse et al., 2013). Since joint quasi-stiffness does not change for different levels of muscle co-contraction, it is not an accurate representation of joint stiffness generated by muscle co-contraction. From another perspective, quasi-stiffness represents response in joint moment due to changes in not only joint position but also muscle activation and joint velocity (Sartori et al., 2015). To address these concerns, joint stiffness is defined in the present study as the elastic response to only changes in joint position. This definition follows the recommendation of Latash and Zatsiorsky (1993) and provides a reasonable basis for the evaluation of the relationship between muscle co-contraction and joint stiffness.

The Co-Contraction Index (CCI) is a commonly used method for quantifying muscle co-contraction. Two common variants of CCI (Falconer and Winter, 1985; Rudolph et al., 2000) allow clinical researchers to make a fast and easy assessment of muscle co-contraction using surface electromyography (EMG) data. EMG-based CCIs have been used to quantify muscle co-contraction in a number of studies (Kellis, 1998; Di Nardo et al., 2015; Banks et al., 2017). However, the more representative method to quantify muscle co-contraction is through estimating individual muscle forces (Kellis et al., 2003), which may require advanced capabilities in neuromusculoskeletal modeling and simulation. Simulation-based studies can estimate muscle forces and compute CCI with either muscle forces (Trepczynski et al., 2018) or the joint moments generated by individual muscles (Souissi et al., 2018). Since CCI is commonly used by the clinical community, it can be used to investigate the relationship between muscle co-contraction and joint stiffness during gait.

Computation of CCI involves choosing from a wide selection of methods. Previous studies have examined how differences in method affect the results of CCI (Knarr et al., 2012; Banks et al., 2017; Souissi et al., 2017). Knarr et al. (2012) calculated CCI based on a formulation by Rudolph et al. (2000) using normalized joint moments from an antagonistic pair of muscles, while Souissi et al. (2017) calculated CCI based on a formulation by using the sum of antagonist moments around each joint. Not only would these choices result in varied CCI outcomes, but they also could affect how the relationship between co-contraction and joint stiffness is interpreted. It would, therefore, be valuable to evaluate how different methodological choices for calculating a CCI affect the CCI’s ability to approximate joint stiffness trends.

This study provides a quantitative evaluation of the relationship between lower extremity muscle co-contraction indices and corresponding joint stiffness during gait for an individual post-stroke. A total of 14 different classes of CCIs were investigated and their correlation with joint stiffness calculated by a personalized neuromusculoskeletal model was assessed. The 14 different classes of CCIs were calculated applying two commonly used formulations (Falconer and Winter, 1985; Rudolph et al., 2000), and each with 7 types of quantities used to represent muscle co-contraction: 4 variations of EMG signals; 2 types of joint moment generated by individual muscles, normalized or not; and the sum of joint moment generated by antagonistic muscles. For CCIs computed from individual pairs of antagonistic muscles, all possible combinations of eligible muscles were covered (12 for the hip joint, 16 for the knee joint, and 3 for the ankle joint). Based on the correlation observed between different CCI calculation methods and joint stiffness, we provide recommendations for the CCI calculation methods that best approximate joint stiffness trends. These recommendations are tailored to fit clinical practice and different levels of capability in neuromusculoskeletal modeling and simulation.

## 2 Methods

### 2.1 Experimental data

Ten cycles of previously published walking data collected from a hemiparetic male subject post-stroke (Meyer et al., 2017) were used for this study. The subject (height 1.70 m, mass 80.5 kg, age 79 years) walked on a split-belt instrumented treadmill (Bertec Corp., Columbus, OH) at a self-selected speed of 0.5 m/s for multiple gait cycles, while motion capture (Vicon Corp., Oxford, UK), ground reaction (Bertec Corp., Columbus, OH) and EMG data (Motion Lab Systems, Baton Rouge, LA) were collected. EMG signals were collected from 16 muscles in each leg (Table 1) using a combination of surface and indwelling electrodes at a frequency of 1000 Hz (Meyer et al., 2017). For more details about the data collection and the experimental protocol, see Meyer et al. (2017).

**Table 1:** Muscles analyzed in this study. The functional role of each muscle is indicated as FLEX – flexor, EXT – extensor, DF – dorsi-flexor, PF – plantar-flexor. A shaded cell for functional role indicates muscle has switched role during gait. EMG scale factor is the scale factor applied to the basic EMG signal of a muscle to account for the difference between true maximum and maximum observed experimentally.

| Muscle Name and Abbreviation | EMG Source | Functional Role |  |  | EMG Scale |  |
| --- | --- | --- | --- | --- | --- | --- |
|  |  | Hip | Knee | Ankle | L | R |
| adductor brevis (addbrev) | adductor longus | FLEX |  |  | 0.14 | 0.05 |
| adductor longus (addlong) |  | FLEX |  |  | 0.33 | 0.07 |
| adductor magnus distal (addDist) |  | EXT |  |  | 0.07 | 0.05 |
| adductor magnus ischial (addmagIsch) |  | EXT |  |  | 0.05 | 0.05 |
| adductor magnus middle (addmagMid) |  |  |  |  | 0.07 | 0.05 |
| adductor magnus proximal (addmagProx) |  | FLEX |  |  | 0.13 | 0.05 |
| gluteus maximus superior (glmax1) | gluteus maximus | EXT |  |  | 0.32 | 0.20 |
| gluteus maximus middle (glmax2) |  | EXT |  |  | 0.33 | 0.20 |
| gluteus maximus inferior (glmax3) |  | EXT |  |  | 0.32 | 0.20 |
| gluteus medius anterior (glmed1) | gluteus medius | EXT |  |  | 0.71 | 0.62 |
| gluteus medius middle (glmed2) |  | EXT |  |  | 0.69 | 0.63 |
| gluteus medius posterior (glmed3) |  | EXT |  |  | 0.69 | 0.62 |
| gluteus minimus anterior (glmin1) |  |  |  |  | 0.32 | 0.15 |
| gluteus minimus middle (glmin2) |  |  |  |  | 0.30 | 0.16 |
| gluteus minimus posterior (glmin3) |  | EXT |  |  | 0.30 | 0.15 |
| iliacus (iliacus) | iliopsoas * | FLEX |  |  | 0.05 | 0.05 |
| psoas (psoas) |  | FLEX |  |  | 0.99 | 0.82 |
| semimembranosus (semimem) | semimem | EXT | FLEX |  | 0.34 | 0.40 |
| semitendinosus (semiten) |  | EXT | FLEX |  | 0.30 | 0.40 |
| biceps femoris long head (bflh) | bflh | EXT | FLEX |  | 0.76 | 0.38 |
| biceps femoris short head (bfsh) |  |  | FLEX |  | 0.76 | 0.39 |
| rectus femoris (recfem) | rectus femoris | FLEX | EXT |  | 0.48 | 0.28 |
| vastus medialis (vasmed) | vastus medialis |  | EXT |  | 0.27 | 0.50 |
| vastus intermedius (vasint) |  |  | EXT |  | 0.31 | 0.44 |
| vastus lateralis (vaslat) | vastus lateralis |  | EXT |  | 0.32 | 0.11 |
| lateral gastronemius (gaslat) | gasmed |  | FLEX | PF | 0.05 | 0.12 |
| medial gastronemius (gasmed) |  |  | FLEX | PF | 0.14 | 0.14 |
| tibialis anterior (tibant) | tibialis anterior |  |  | DF | 0.61 | 1.00 |
| tibialis posterior (tibpost) | tibialis posterior * |  |  | PF | 0.05 | 0.05 |
| peroneus brevis (perbrev) | peroneus longus |  |  | PF | 0.05 | 0.98 |
| peroneus longus (perlong) |  |  |  | PF | 0.05 | 0.99 |
| peroneus tertius (pertert) |  |  |  | DF | 0.05 | 0.99 |
| soleus (soleus) | soleus |  |  | PF | 0.65 | 0.96 |
| extensor digitorum longus (edl) | edl * |  |  | DF | 1.00 | 0.26 |
| flexor digitorum longus (fdl) | fdl * |  |  | PF | 1.00 | 0.05 |

\* measured using indwelling EMG

### 2.2 EMG data processing and EMG-driven model calibration

Collected EMG data were high-pass filtered at 40 Hz, demeaned, rectified, and low-pass filtered at a variable cutoff frequency of 3.5/period of the gait cycle being processed (Lloyd and Besier, 2003) while using a 4^th^ order zero phase-lag Butterworth filter (Meyer et al., 2017). Filtered EMG data were subsequently normalized to the maximum value over all trials and resampled to 101 normalized time points for each gait cycle. Normalized EMG data for each gait cycle were offset by the minimum value so that the minimum EMG value for each gait cycle was zero. These processing procedures represent the basic approach for processing EMG adopted by other studies for the quantification of muscle co-contraction (Rosa et al., 2014). The EMG signals processed under the aforementioned procedures were defined as the basic EMG signals, *EMG*_*basic*_.

An EMG-driven model was used to calibrate parameters of the model, including a few for conditioning basic EMG to muscle excitation along with those for musculotendon force functions, within an optimization process to match the joint moments produced by the muscles in the model with those calculated from inverse dynamics (Meyer et al., 2017). The conditioning of the EMG signals involved adding an electromechanical delay and applying a muscle-specific EMG scale factor to *EMG*_*basic*_. The electromechanical delay is defined as the duration from the instant an electrical signal is received to the instant a force response is generated by the muscle. The electromechanical delay was assumed to be fixed for all the muscles in each leg (Meyer et al., 2017), which was determined by the optimization to be 82 ms and 93 ms for the left and right side, respectively. The muscle-specific scale factor (Table 1) was used to account for the difference between the true maximum EMG value and maximum value over all experimental trials. The EMG signals resulted from calibration for both electromechanical delay and scaling were defined as fully calibrated EMG signals, *EMG*_*calibrated*_.

To isolate the underlying effect of the two EMG parameters on quantification of co-contraction, we introduced two additional types of EMG signals: 1) scaled EMG signals *EMG*_*scaled*_, which are EMG signals normalized to the optimized maximum EMG value but without electromechanical delay; 2) delayed EMG signals *EMG*_*delayed*_, which are electromechanically delayed EMG signals but not normalized to the optimized maximum value. These signals were obtained by

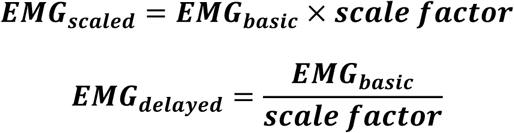

The EMG-driven model calibration process also provided estimates of force and joint moment(s) generated by each muscle during walking. The muscle force and joint moment were passed onto CCI computation and joint stiffness estimation.

### 2.3 CCI computation

CCI values were computed from the two commonly used formulations. *CCI*_*1*_ was based on the study by Rudolph et al. (2000):

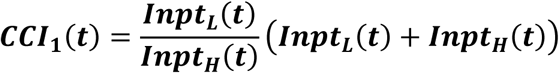

*CCI*_*2*_ was based on the work by Falconer and Winter (1985):

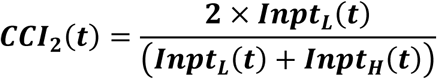

For both formulations, the lower input *Inpt*_*L*_ represents input quantity with lower absolute magnitude, and higher input *Inpt*_*H*_ represents input quantity with higher absolute magnitude sampled at time instant *t*, which is each of the 101 normalized time points (0 – 100% of gait cycle at 1% increment) of a gait cycle. For EMG-based CCI, the input quantities include the aforementioned four types of EMG signals (*EMG*_*basic*_, *EMG*_*scaled*_, *EMG*_*delayed*_, and *EMG*_*calibrated*_). For moment-based CCI, the input quantities include joint moment from an antagonistic pair of muscles, actual values (*M*_*mus*_), or values normalized to maximum values over all selected trials (*M*_*mus norm*_) and the sum of antagonist moment around the joints *M*_*sum*_. This study uses a notation to specify the formulation and input quantities used to compute CCI, e.g., *EMG*_*delayed*_-based *CCI*_*1*_ means the CCI values are calculated using delayed EMG signals based on Rudolph et al. (2000) formulation.

*CCI*_*1*_ can also be seen as the product of two terms where one term represents the ratio between antagonist muscle activities while the other represents the sum of antagonist muscle activities:

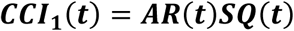

where antagonist ratio *AR* represents the ratio of the antagonist quantities used for CCI computation and sum of antagonist quantities for CCI calculation *SQ* represents the sum of the antagonist quantities:

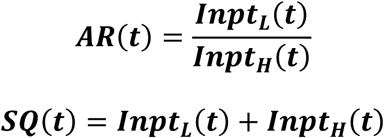

In addition to varying the types of quantities used to compute CCI, this study also investigated how the difference in constituent muscles for an antagonistic pair could affect the relationship between CCI and joint stiffness. CCI was computed for three lower limb degrees of freedom (DOFs) in the sagittal plane: hip, knee, and ankle flexion and extension for both non-paretic side and paretic side. Lower extremity muscles were classified by their functional roles during gait (Table 1), and one muscle was selected from each of the agonist group and the antagonist group to form various combinations of antagonistic pairs. Antagonistic muscle pairs consisting of small muscles that were not major contributors to overall joint stiffness (less than 2% on average) were not included for the subsequent analyses. The majority of the EMG-based CCI in previous studies was computed using EMG signals measured from surface muscles (Rosa et al., 2014) because the alternative indwelling EMG method is invasive and not universally available. Therefore, despite the availability of indwelling EMG data of deep muscles (iliacus, psoas, tibialis posterior, extensor, and flexor digitorum longus), the analyses in this study focused on CCI computed from surface EMG signals of muscles for the findings to be more applicable in clinical settings. Antagonistic muscle pairs consisting of the aforementioned deep muscles were not included for the subsequent analyses either. After removing antagonistic pairs consisted of small or deep muscles from consideration, CCI values from 12 antagonistic muscle pairs for hip joint, 16 for knee joint, and 3 for ankle joint were analyzed.

### 2.4 Estimation of joint stiffness

Sagittal plane stiffness of the lower extremity joints (hip, knee, and ankle) of the subject was estimated using a model-based formulation (Shourijeh and Fregly, 2020):

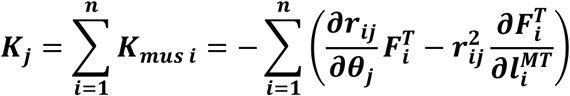

where stiffness *K*_*j*_ of DOF *j* is equal to the sum of the individual muscle stiffnesses *K*_*mus i*_. *r*_*ij*_ represents the moment arm of the *i*^*th*^ muscle about DOF *j. θ*_*j*_ is the joint angle about DOF *j. F*^*T*^_*i*_ and *l*^*MT*^_*i*_ represent the tendon force and muscle-tendon length of muscle *i*. The model-based stiffness formulation assumes a rigid-tendon muscle model. As moment arms and muscle-tendon lengths of the musculoskeletal model (Meyer et al., 2017) were represented by surrogate models in the form of polynomials of joint kinematics, muscle stiffness around a joint could be computed analytically. Joint stiffness was estimated at the same % points of gait at which CCI was evaluated.

### 2.5 Statistical Analyses

The strength of association between CCI (*CCI*_*1*_ or *CCI*_*2*_) and joint stiffness (*K*_*joint*_) was determined by the Pearson correlation coefficients using the *corrcoef* function in MATLAB (MathWorks, Natick, USA) between the two time series for each of the 10 gait cycles analyzed.

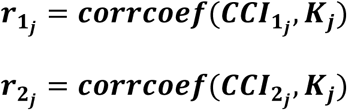

Since *AR* and *SQ* were available, the correlation between these *CCI*_*1*_ components and joint stiffness was also investigated to elucidate which component was the main driver for correlation *r*_*l*_.

The Wilcoxon rank sum test implemented in MATLAB as the *Ranksum* function was used to compare the mean correlation coefficient between the two classes of data (10 correlation coefficients per class for 10 gait cycles). The analysis tested the null hypothesis that the two classes of data came from samples with continuous distributions possessing equal medians.

## 3 Results

A consistency across trials was observed in the estimates of joint stiffness at all three joints (Figs. 1A and 1B, 1^st^ row). For the hip joint on each side, joint stiffness continued to increase in the early in the stance phase (0 – 15% gait cycle), maintained at the level for the remainder of the stance phase (15% - 55% gait cycle) and continued to decrease during late stance and swing phases (55% – 100% gait cycle). For the knee joint on each side, joint stiffness continued to increase early in the stance phase (0 – 20% gait cycle) and began to gradually decrease from the peak value. For the ankle joint on the non-paretic side, joint stiffness continued to rise during the stance phase (0 – 60%) before descending and on the paretic side, joint stiffness continued to increase in the early in the stance phase (0 – 20% gait cycle), maintained at the level for the remainder of the stance phase (20% - 65% gait cycle) and continued to decrease during swing phases (65% – 100% gait cycle). Examples of CCI results were also plotted to visualize the correlation between joint stiffness and CCI (Figs. 1A and 1B, 2^nd^ row: EMG-based *CCI*_1_, 3^rd^ row: moment-based *CCI*_1_, 4^th^ row: EMG-based *CCI*_2_ and 5^th^ row: moment-based *CCI*_2_).

**Figure 1:**
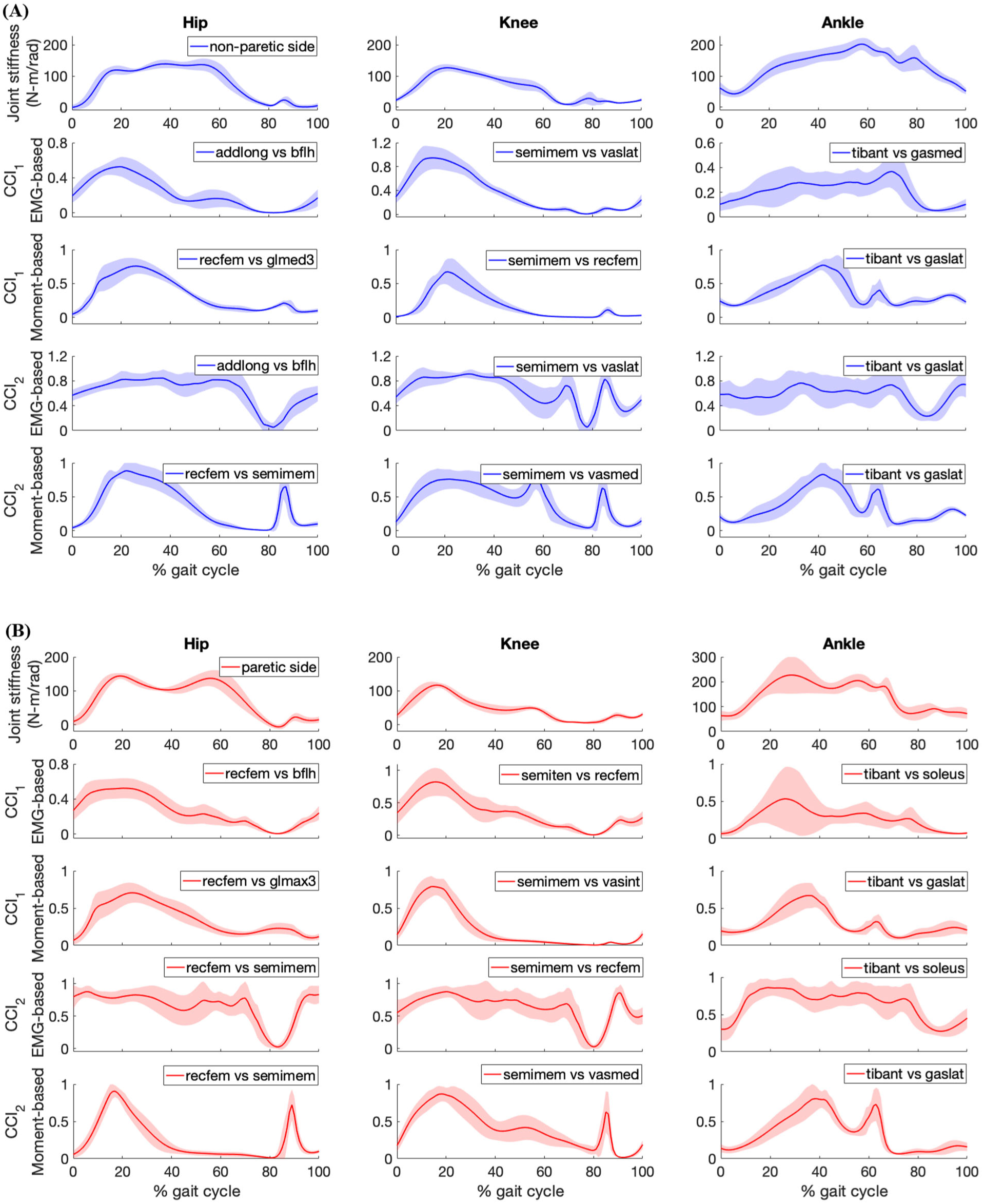
Stiffness, sample *CCI*_1_ and *CCI*_2_ values (mean ± 1 standard deviation) around lower extremity joints of the subject during gait for (A) non-paretic side data in blue, and (B) paretic side data in red. The antagonistic pair of muscles selected for CCI computation are identified in the legend of the subplot. EMG-based CCI shown in the figure is computed using *EMG*_*delayed*_ signals and moment-based CCI are computed using unnormalized joint moment and then normalized by the maximum value over all gait trials analyzed.

Correlation r_1_ between the values of *CCI*_1_ and joint stiffness were moderate (0.5 < *r*_*1*_ < 0.7, Moore et al., 2015) for the hip joint, strong (*r*_*1*_ > 0.7) for the knee joint and moderate (0.5 < *r*_*1*_ < 0.7) for the ankle joint (Fig. 2). The highest mean values of *r*_*1*_ for EMG-based and moment-based *CCI*_*1*_ were 0.51 and 0.70, respectively at the hip joint on the non-paretic side; 0.59 and 0.66 at the hip joint on the paretic side; 0.81 and 0.85 at the knee joint on the non-paretic side; 0.91 and 0.91 at the knee joint on the paretic side; 0.57 and 0.38 at the ankle joint on the non-paretic side; 0.76 and 0.59 at the ankle joint on the paretic side.

**Figure 2:**
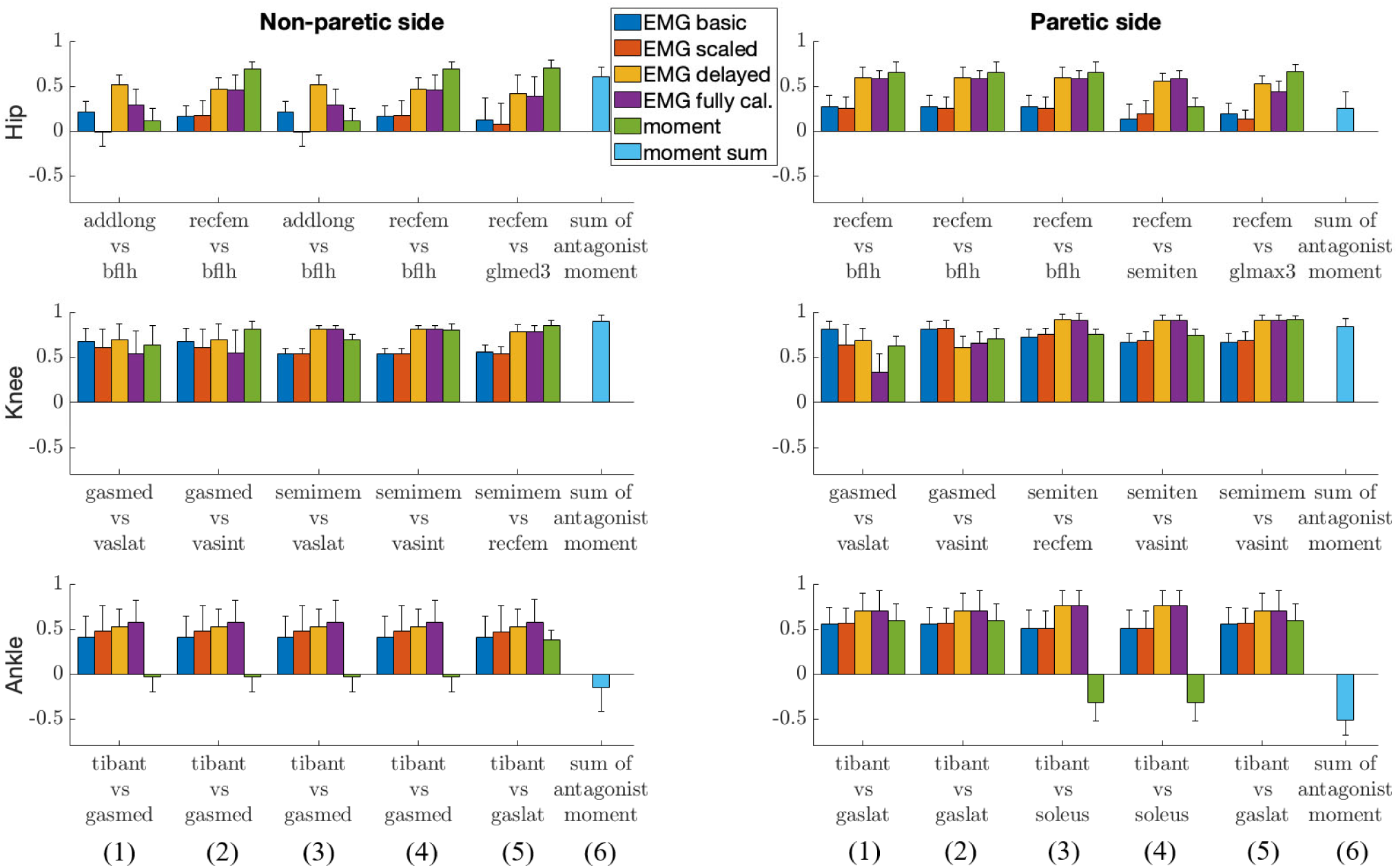
Correlation between *CCI*_1_ and joint stiffness *K*_*joint*_. Bars are at the mean value of the Pearson correlation coefficient, and error bars are at 1 standard deviation (+/– depending on the sign of mean value). Each muscle combination for antagonistic pairing displayed in the figure represents the best-in-class correlation between *K*_*joint*_ and *CCI*_*1*_ computed using a specific type of quantity: (1) *EMG*_*basic*_ (in blue); (2) *EMG*_*scaled*_ (in red); (3) *EMG*_*delayed*_ (in orange); (4) *EMG*_*calibrated*_ (in purple); (5) *M*_*mus*_ (in green). Bar (6) represents the correlation between *K*_*joint*_ and *CCI*_*1*_ computed using the sum of the antagonist moment around each joint, *M*_*sum*_ (in cyan).

Correlation r_2_ between the values of *CCI*_2_ and joint stiffness were moderate (0.5 < *r*_*2*_ < 0.7) for the hip joint, moderate (0.5 < *r*_*2*_ < 0.7) for the knee joint, weak (0.3 < *r*_*2*_ < 0.5) for the ankle joint on the non-paretic side and moderate (0.5 < *r*_*2*_ < 0.7) on the paretic side (Fig. 3). The highest mean value of *r*_*2*_ for EMG-based and moment-based *CCI*_*2*_ were 0.69 and 0.64 respectively at the hip joint on the non-paretic side; 0.54 and 0.38 at the hip joint on the paretic side; 0.64 and 0.79 at knee joint on the non-paretic side; 0.60 and 0.79 at the knee joint on the paretic side; 0.40 and 0.42 at the ankle joint on non-paretic side; 0.59 and 0.62 at the ankle joint on the paretic side.

**Figure 3:**
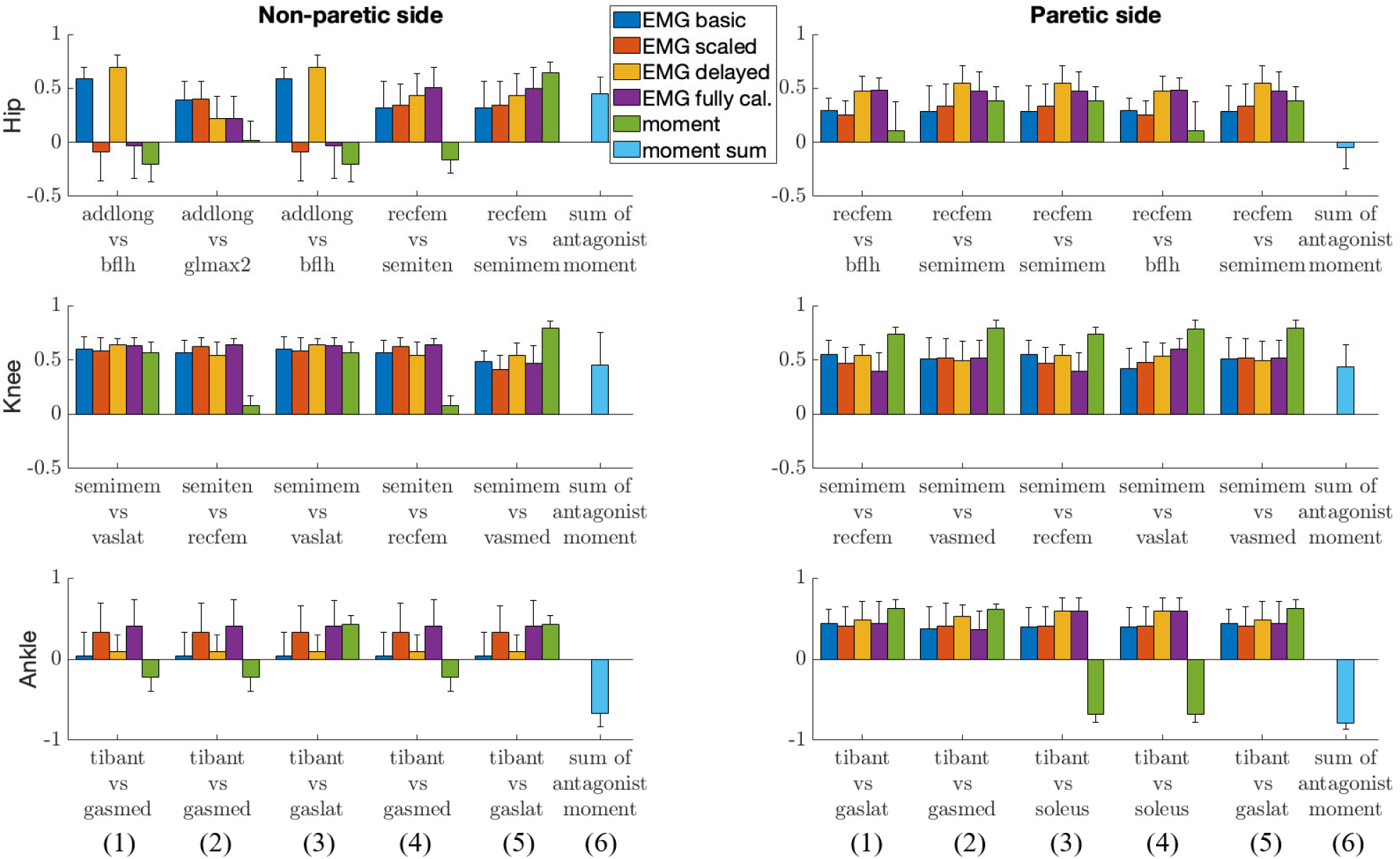
Correlation between *CCI*_2_ and joint stiffness *K*_*joint*_. Bars are at the mean value of the Pearson correlation coefficient and error bars are at 1 standard deviation (+/– depend on the sign of mean value). Each muscle combination for antagonistic pairing displayed in the figure represents the best-in-class correlation between *K*_*joint*_ and *CCI*_*2*_ computed using a specific type of quantity: (1) *EMG*_*basic*_ (in blue); (2) *EMG*_*scaled*_ (in red); (3) *EMG*_*delayed*_ (in orange); (4) *EMG*_*calibrated*_ (in purple); (5) *M*_*mus*_ (in green). Bar (6) represents the correlation between *K*_*joint*_ and *CCI*_*2*_ computed using the sum of the antagonist moment around each joint, *M*_*sum*_ (in cyan).

Differences in the mean between best-in-class correlations indicated that *r*_*1*_ was higher than *r*_*2*_ in most cases (Table 2). For *EMG*_*delayed*_ and *EMG*_*calibrated*_ cases at the hip (non-paretic side) and *M*_*mus*_ class at the ankle (both sides), best-in-class mean values of *r*_*2*_ were higher than those of *r*_*1*_.

**Table 2:**
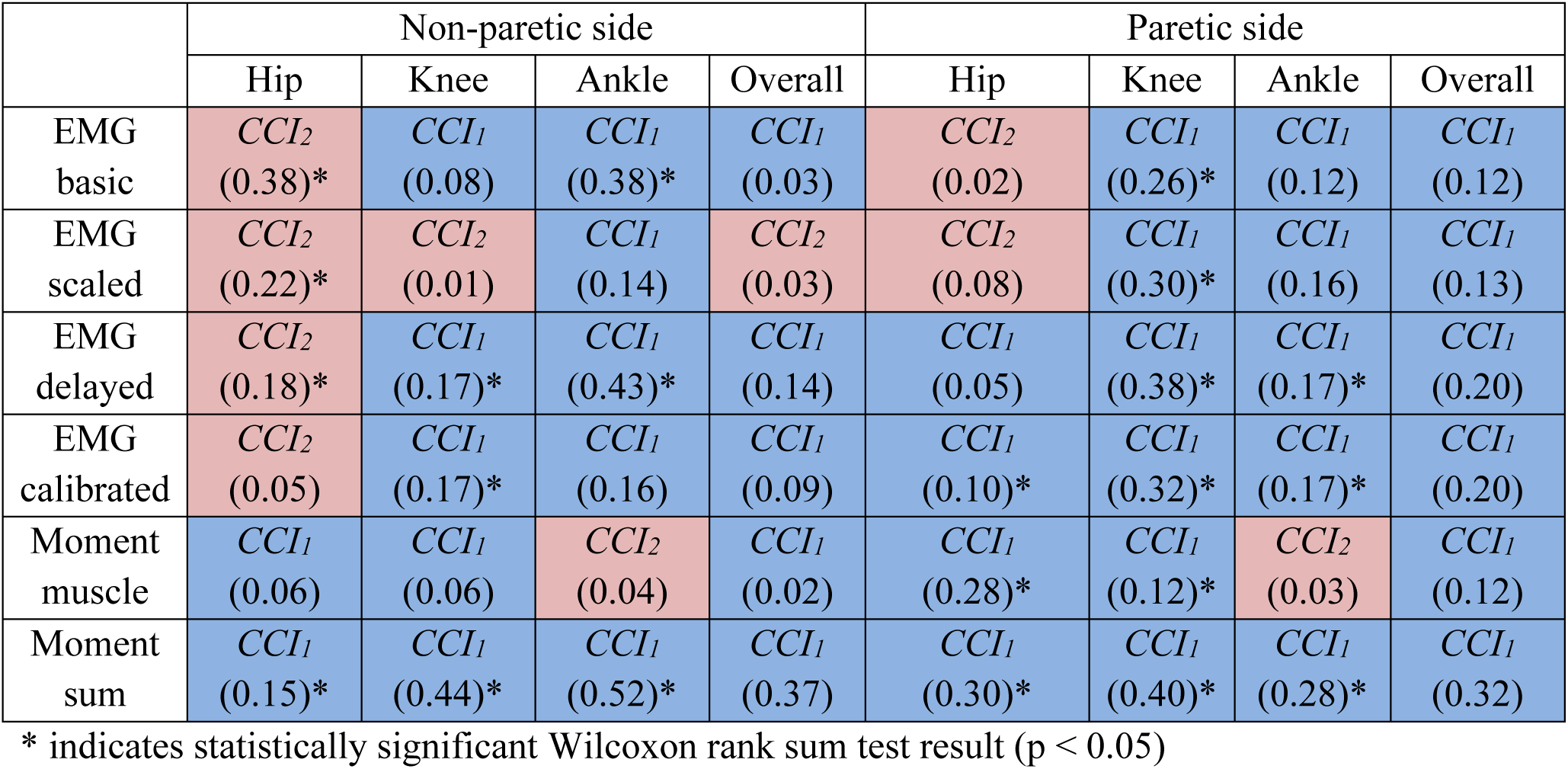
Mean difference in the highest correlation for each case between both CCI methods. The method which produces a higher correlation with *K*_*joint*_ for each case is identified in the corresponding table cells. Blue shading indicates mean correlation *r*_*1*_ is higher than r_*2*_, whereas red shade indicates the mean correlation *r*_*2*_ is higher than r_*1*_. Each value in the Overall column is the mean of values from the corresponding Hip, Knee, and Ankle column.

|  | Non-paretic side |  |  |  | Paretic side |  |  |  |
| --- | --- | --- | --- | --- | --- | --- | --- | --- |
|  | Hip | Knee | Ankle | Overall | Hip | Knee | Ankle | Overall |
| EMG basic | $CCI_2$<br>(0.38)* | $CCI_1$<br>(0.08) | $CCI_1$<br>(0.38)* | $CCI_1$<br>(0.03) | $CCI_2$<br>(0.02) | $CCI_1$<br>(0.26)* | $CCI_1$<br>(0.12) | $CCI_1$<br>(0.12) |
| EMG scaled | $CCI_2$<br>(0.22)* | $CCI_2$<br>(0.01) | $CCI_1$<br>(0.14) | $CCI_2$<br>(0.03) | $CCI_2$<br>(0.08) | $CCI_1$<br>(0.30)* | $CCI_1$<br>(0.16) | $CCI_1$<br>(0.13) |
| EMG delayed | $CCI_2$<br>(0.18)* | $CCI_1$<br>(0.17)* | $CCI_1$<br>(0.43)* | $CCI_1$<br>(0.14) | $CCI_1$<br>(0.05) | $CCI_1$<br>(0.38)* | $CCI_1$<br>(0.17)* | $CCI_1$<br>(0.20) |
| EMG calibrated | $CCI_2$<br>(0.05) | $CCI_1$<br>(0.17)* | $CCI_1$<br>(0.16) | $CCI_1$<br>(0.09) | $CCI_1$<br>(0.10)* | $CCI_1$<br>(0.32)* | $CCI_1$<br>(0.17)* | $CCI_1$<br>(0.20) |
| Moment muscle | $CCI_1$<br>(0.06) | $CCI_1$<br>(0.06) | $CCI_2$<br>(0.04) | $CCI_1$<br>(0.02) | $CCI_1$<br>(0.28)* | $CCI_1$<br>(0.12)* | $CCI_2$<br>(0.03) | $CCI_1$<br>(0.12) |
| Moment sum | $CCI_1$<br>(0.15)* | $CCI_1$<br>(0.44)* | $CCI_1$<br>(0.52)* | $CCI_1$<br>(0.37) | $CCI_1$<br>(0.30)* | $CCI_1$<br>(0.40)* | $CCI_1$<br>(0.28)* | $CCI_1$<br>(0.32) |

\* indicates statistically significant Wilcoxon rank sum test result ( $p < 0.05$ )

Correlation between SQ and joint stiffness was generally stronger than that between AR and joint stiffness (Fig 4), suggesting the association between *CCI*_*1*_ and joint stiffness is driven more by the SQ component. Correlation between *AR* and joint stiffness in some cases was in the range from negative to low positive, yet the correlation between *CCI*_*1*_ and joint stiffness was at the moderate-to-strong level because of the high correlation between *SQ* and joint stiffness.

**Figure 4:**
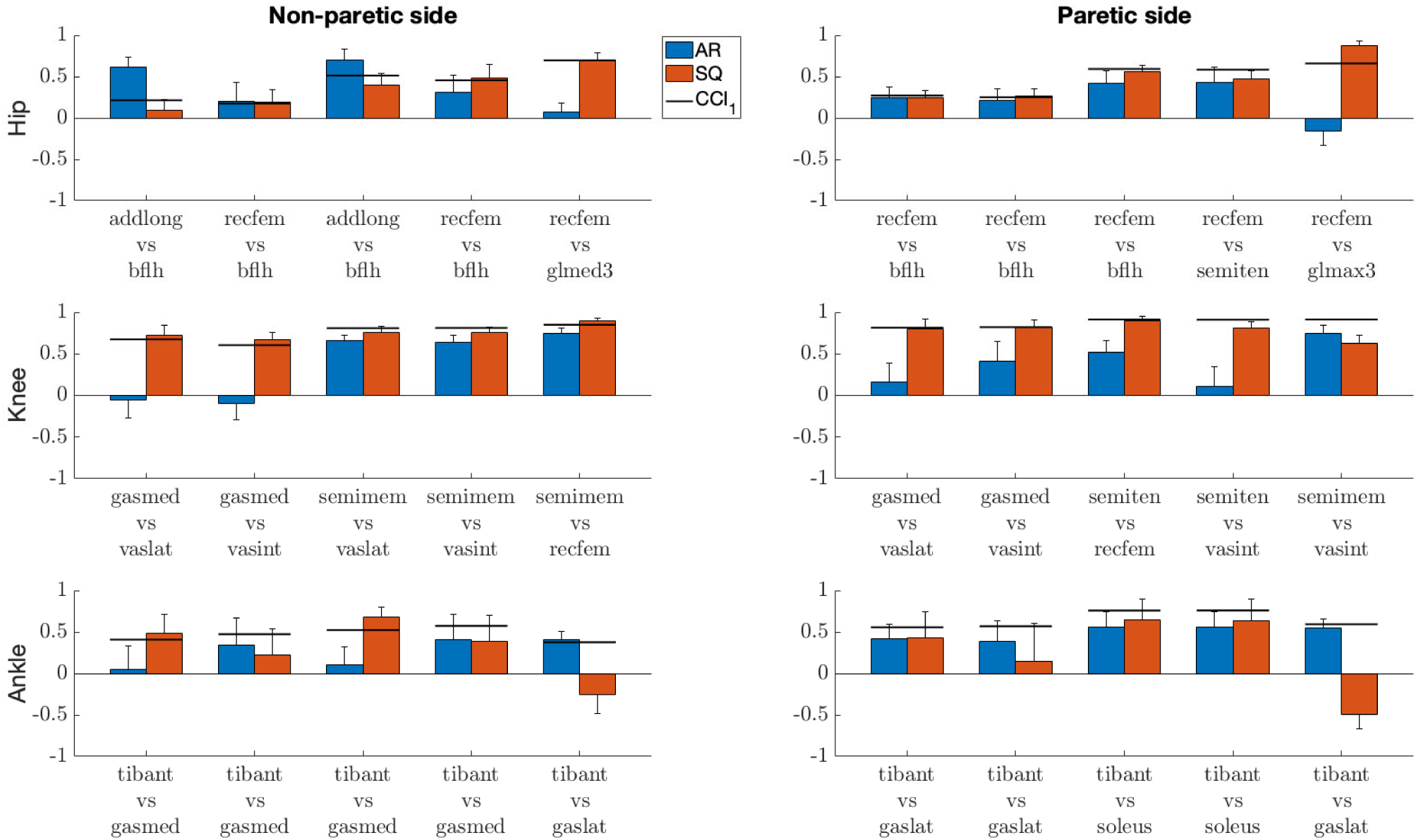
Correlation between *CCI*_*1*_ terms (*AR* and *SQ*) and joint stiffness. Bars are at the mean value of the Pearson correlation coefficient, and error bars are at 1 standard deviation (+/– depending on the sign of mean value). Blue bars are for *AR*, and red bars are for *SQ*. Dark lines are at the mean value of the correlation coefficient between *CCI*_*1*_ and joint stiffness.

## 4 Discussion

This study evaluated how well different CCI formulations approximate lower extremity joint stiffness trends during gait. Although moderate to strong correlation was observed between some CCI formulations and corresponding joint stiffness, this correlation was highly dependent on the methodological choices made for CCI computation. The following observations summarize the conditions under which high CCI correlations with joint stiffness were obtained: (1) *CCI*_*1*_ formulation (Rudolph et al., 2000) was more correlated than *CCI*_*2*_ formulation (Falconer and Winter, 1985); (2) Moment-based *CCI*_*1*_ from individual antagonistic muscle pairs was more correlated than EMG-based *CCI*_*1*_; (3) *M*_*mus*_ and *EMG*_*delayed*_ were the most correlated for moment-based *CCI*_*1*_ and EMG-based *CCI*_*1*_, respectively. These findings could be used as preliminary foundation for predicting joint stiffness from EMG-based measurement of muscle co-contraction.

Correlation between *CCI*_*1*_ and joint stiffness was higher than for *CCI*_*2*_ for most cases (Table 2), whether CCI was EMG-based or moment-based. Joint stiffness *K*_*joint*_ is a sum of the stiffness generated by all the individual muscles *K*_*mus*_. The *SQ* term in the *CCI*_*1*_ formulation was more effective at characterizing this summation than any term in the *CCI*_*2*_ formulation. This effectiveness became more pronounced when the quantities used for CCI computation from each muscle were accurate proxies for the corresponding *K*_*mus*_. On the other hand, the *CCI*_*2*_ formulation was more suitable for quantifying the ratio of quantities from two antagonist muscles. This observation is demonstrated by the high correlation between results from *CCI*_*2*_ and those from the *AR* term in *CCI*_*1*_. Close examination also showed that *CCI*_*2*_ values reached a peak in magnitude in the swing phase (∼60 to 100% gait cycle) comparable to that during the stance phase (0% to ∼60% gait cycle) at several joints (Fig. 1A and 1B). This phenomenon was deemed unlikely to be physiological. This exposes the limitation of the *CCI*_*2*_ formulation that when the two quantities from antagonist muscles become close in magnitude, *CCI*_*2*_ would report a high level of co-contraction regardless of how small both quantities might be, as *CCI*_*2*_ focuses on quantifying the ratio between the two quantities. Since the *CCI*_*1*_ formulation was a better choice than *CCI*_*2*_ for approximating joint stiffness trends, the subsequent discussion will focus on the methodological choices involved in calculating *CCI*_*1*_.

For each joint, the best correlation between *M*_*mus*_-based *CCI*_*1*_ results and joint stiffness was generally higher than the best correlation between any EMG-based *CCI*_*1*_ results and joint stiffness. The higher correlation between *M*_*mus*_-based *CCI*_*1*_ and joint stiffness was because the joint moments produced by individual muscles could serve as a more accurate proxy for *K*_*mus*_ than could EMG signals. Additional analyses revealed that 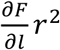 was the dominant term in *K*_*mus*_ (responsible for > 95% of *K*_*mus*_ magnitude) for the vast majority of the muscles included in the model. The 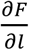 term can be further expressed as the product of muscle force *F*_*mus*_ and a complex function consisting of geometric and dynamic muscle parameters. *M*_*mus*_ is the product of moment arm *r* and muscle force *F*_*mus*_, both being present in the 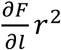 term. In contrast, EMG amplitude partially reflects *F*_*mus*_. However, it is well known that the relationship between EMG amplitude and *F*_*mus*_ is not linear, especially when large force is generated (Buchanan et al., 2004). As a result, the joint moment generated by a muscle carries more information about joint stiffness generation than does the EMG signal of the muscle. Previous studies concluded that moment-based CCI offered a more complete description of muscle activity and joint loads than does EMG-based CCI (Knarr et al., 2012; Souissi et al., 2017). This study built upon this conclusion and found that moment-based *CCI*_*1*_ also provided better ability at approximating joint stiffness trends than the EMG-based *CCI*_*1*_.

Among the moment-based *CCI*_*1*_ methods, the ones based on *M*_*mus*_ were most likely to achieve the highest correlation with joint stiffness as compared to the ones based on either *M*_*sum*_ or *M*_*mus_norm*_. The majority of the muscles analyzed articulate more than one DOF. Some muscles articulate DOFs that were not analyzed in this study, e.g. hip adduction-abduction and ankle inversion-eversion. Some muscles even switch their roles during dynamic tasks. For instance, although psoas is primarily a hip flexor, it has a minor ab/adduction role in our musculoskeletal model and switches between being a hip adductor and a hip abductor during gait. By summing the moments, we could not separate the collective muscle activities and allocate the precise proportion of muscle activities of each joint to account for stiffness generation. Therefore, the *M*_*sum*_-based *CCI*_*1*_ correlated poorly with joint stiffness for joints with multiple DOFs (i.e., the hip and ankle) but correlated well with joint stiffness for joints with only a single DOF (i.e., the knee joint). A previous comparison study by Knarr et al. (2012) on CCI methods used the *M*_*mus_norm*_ value to compute a CCI. In our study, for most antagonistic pairs analyzed, the *M*_*mus norm*_-based *CCI*_*1*_ had a lower correlation with joint stiffness than did the *M*_*mus*_-based *CCI*_*1*_ because of a lower correlation between the *SQ* term and joint stiffness for *M*_*mus_norm*_-based *CCI*_*1*_ than *M*_*mus*_-based *CCI*_*1*_. As Souissi et al. (2017) argued, the use of a normalized moment did not allow distinguishing between the force-producing capacity of the agonist and antagonist muscle. Therefore, the relative contribution of each quantity to the *SQ* term was distorted by the normalization. In contrast, computation of the *M*_*mus*_-based *CCI*_*1*_ could potentially include two antagonistic muscles that the sum of the joint moment generated by these muscles would correlate well with joint stiffness. In other words, the *SQ* term in *CCI*_*1*_, which is equal to the sum of the *M*_*mus*_ quantities, would help *CCI*_*1*_ to also correlate well with joint stiffness. However, to obtain reliable *M*_*mus*_ data, one must use a calibrated musculoskeletal model. Consequently, the moment-based CCI is less frequently adopted by the clinical community than the more accessible alternatives, e.g. the EMG-based CCI.

For a clinical setting, the simplest and easiest approach is to use a CCI that requires only EMG data. However, EMG processing methods affect the correlation between EMG-based CCIs and joint stiffness. From the *EMG*_*basic*_ signals, two modifications were applied to obtain the other types of processed EMG signals. One modification was adding electromechanical delay from *EMG*_*basic*_ to *EMG*_*delayed*_. The delays were set to 82 and 93 ms for the non-paretic side and paretic side respectively, based on the optimal values found via the EMG-driven model. This modification increased the correlation between *CCI*_*1*_ and *K*_*joint*_. As previously discussed, EMG signals are able to convey partial information about joint stiffness because of the relationship between EMG amplitude and *F*_*mus*_. Introducing electro-mechanical delay improves the synchronization between an EMG signal the resulting muscle force. Consequently, the correlation between the EMG signal and *K*_*mus*_ increases, resulting in an increased correlation between the *EMG*_*delayed*_-based *CCI*_*1*_ and *K*_*joint*_. The second modification was applying a muscle-specific scale factor (Table 2), i.e. from *EMG*_*basic*_ to *EMG*_*scaled*_ and from *EMG*_*delayed*_ to *EMG*_*calibrated*_. This modification did not produce a clear improvement in the correlation between resultant *CCI*_*1*_ and *K*_*joint*_. Applying the scale factor did not change the ability of the EMG signals to represent *K*_*mus*_, and the correlation between scaled EMG signal and *K*_*mus*_ remained the same as before scaling. However, the muscle-specific scale factor did change the relative contribution of muscle EMG amplitudes to the *SQ* term in the *CCI*_*1*_ formulation. In some cases, a change in *SQ* caused a decrease in the correlation between resultant *CCI*_*1*_ and *K*_*joint*_. Although applying muscle-specific scale factors changed muscle force estimates during the calibration of the EMG-driven musculoskeletal model, these scale factors did not consistently increase the correlation between *CCI*_*1*_ and *K*_*joint*_. Excluding the uncertain effect of scaling and including the improvement from adding electromechanical delay, *EMG*_*delayed*_-based *CCI*_*1*_ appeared to be the most suitable choice for the purpose of approximating joint stiffness trends.

This study presented CCI data from muscles from which EMG data could be obtained by surface measurement. Although indwelling EMG data for some muscles were also collected from the subject studied, these muscles were excluded from the CCI analysis, since EMG data would not be available from them in a clinical setting. Contrary to the general trends noted above, some discrepancies existed: 1) at the hip, the correlation between *CCI*_*2*_ and *K*_*joint*_ was better than that between *CCI*_*1*_ and *K*_*joint*_; 2) at the ankle, the correlation between *M*_*mus*_-based *CCI*_*1*_ and *K*_*joint*_ was lower than between EMG-based *CCI*_*1*_ and *K*_*joint*_. However, when CCI was recalculated by including muscles with available indwelling EMG data, these discrepancies mostly disappeared. Among the muscles omitted were iliacus and psoas, two primary hip flexors. If included on the non-paretic side, the iliacus – bflh pair would become the new leader for correlation between *CCI*_*1*_ and *K*_*joint*_ for the *EMG*_*basic*_, *EMG*_*scaled*_, and *EMG*_*delayed*_ cases and the psoas – bflh pair for the *EMG*_*calibrated*_ case. All held a higher correlation with *K*_*joint*_ than did their *CCI*_*2*_ counterparts. Also among the omitted muscles were edl and tibpost, one primary dorsi-flexor, and one primary plantar-flexor muscle. The mean correlation between *M*_*mus*_-based *CCI*_*1*_ and *K*_*joint*_ for the edl – tibpost pair on the non-paretic side was found to be 0.89, higher than that between any EMG-based *CCI*_*1*_ and *K*_*joint*_ (highest mean = 0.57). Even though indwelling EMG measurements are invasive, the measurement technique seems to provide valuable information on muscle co-contraction and joint stiffness.

One limitation of this study was that the joint stiffness used for comparison with different CCI methods was obtained from a neuromusculoskeletal model instead of experimentally usually a joint perturbation technique. A model-based approach was used since the perturbation approach is difficult to implement experimentally, especially for dynamic tasks such as gait (Pfeifer et al., 2012). The model-based approach has been reported to generate joint stiffness estimates that compare well with experimental joint stiffness measurements for isometric conditions (Pfeifer et al., 2012; Sartori et al., 2015). A published model for estimating joint stiffness (Shourijeh and Fregly, 2020) was used in the present study, and the model parameters were calibrated using a validated EMG-driven modeling process (Meyer et al., 2017). Thus, the model closely reproduced the subject’s experimental joint moments when the subject’s experimental EMG and kinematic data were used as input, suggested that the estimated muscle forces and thus joint stiffness values are likely to be at least reasonable.

Another limitation of this study was that it analyzed gait data collected from only a single hemiparetic subject. Although the collection of sixteen channels (including 6 indwelling) of EMG from each leg of the subject during walking was time-consuming and not a common practice, it facilitated the calibration of our musculoskeletal model. Despite analyzing one subject limits our ability to draw more general conclusions that could be applied to the stroke population, at the same time, this dataset provided a unique opportunity to build a musculoskeletal model of the subject and calibrate the model parameters using an EMG-driven framework that did not require prediction of any missing EMG signals (Sartori et al., 2014). Another benefit of the data set was that the indwelling EMG data allowed us to demonstrate that the use of these data improved CCI correlation with joint stiffness at the hip and ankle.

In conclusion, this study demonstrated the feasibility of using various CCIs to approximate lower limb joint stiffness trends during gait. Numerous methodological choices for CCI computation were examined. Key methodological choices to achieve the highest possible correlation between CCI and joint stiffness should include the use of *CCI*_1_ formulation and the use of *M*_*mus*_ or *EMG*_*delayed*_ as the quantities for computing moment-based or EMG-based *CCI*_*1*_. Antagonistic muscle pairings that yielded the highest correlations between CCI and joint stiffness were also identified. These recommendations on the methodological choices were provided for researchers with different levels of neuromusculoskeletal modeling capability. By using CCI to approximate joint stiffness trends, this study may open an alternative approach to estimate joint stiffness, which is difficult to obtain through either computational modeling or experimental measurement.

## 6 Author Contributions

Conceptualization: GL DA MSS BJF

Funding acquisition: BJF

Data collection: BJF CP

Investigation: GL DA MSS BJF

Methodology: GL DA MSS BJF

Formal analysis: GL DA MSS BJF

Supervision: MSS BJF

Drafting the Manuscript: GL MSS BJF

Revising the Manuscript: GL DA MSS CP BJF

## 7 Acknowledgments

This work was conducted with support from the Cancer Prevention and Research Institute of Texas (CPRIT) funding RR170026. The authors would like to thank Marleny Arones for her assistance with analyzing the data.

## Data Availability Statement

The datasets analyzed for this study are available on request from the corresponding author.

